# Visual Emotion Perception in a Deep Neural Network Model with Both Bottom-Up and Top-Down Connections

**DOI:** 10.64898/2026.04.06.716704

**Authors:** Peng Liu, Ke Bo, Yujun Chen, Andreas Keil, Mingzhou Ding, Ruogu Fang

**Affiliations:** J. Crayton Pruitt Family Department of Biomedical Engineering, Herbert Wertheim College of Engineering, University of Florida, Gainesville, Florida, USA; Department of Psychological and Brain Sciences, Dartmouth College, Hanover, New Hampshire, USA; Center for Cognitive Aging and Memory, McKnight Brain Institute, University of Florida, Gainesville, Florida, USA; Department of Psychology and Center for the Study of Emotion & Attention, University of Florida, Gainesville, Florida, USA

**Keywords:** Emotion perception, top-down feedback, emotion modulation, deep neural networks, NeuroAI, visual cognition, fMRI, amygdala

## Abstract

Emotion reshapes perception by modulating sensory processing through top-down feedback—a process referred to as emotional perception. The computational mechanisms by which distinct affective signals influence visual representations however remain poorly understood. Here, we use a deep neural network to simulate this process and test mechanistic hypotheses about how top-down feedback guides emotional peception. Most existing models treat the perception of emotional content as a static, feedforward task, overlooking the dynamic interplay between internal states, external goals, and sensory input that characterizes affective perception in the brain. We introduce EmoFB, a biologically inspired model that integrates an affective system with a visual processing hierarchy through two functionally distinct feedback signals: intrinsic feedback, arising from the model’s own affective appraisal of perceptual input, and external steering, conveying contextual priors such as task expectations or target categories. We evaluated EmoFB on three tasks varying in perceptual ambiguity (Single Image, Side-by-Side, and Overlay). External steering exerted the strongest influence, not only improving recognition under challenging conditions but also restructuring internal representations by sharpening category-specific clustering in feature space. Crucially, top-down feedback increased brain–model representational similarity, strengthening alignment with human fMRI responses across early visual cortex, ventral visual areas, and the amygdala. EmoFB provides a computational framework for testing neurocognitive theories of emotion appraisal and top-down feedback modulation. It bridges affective neuroscience and artificial intelligence, offering mechanistic insight into how emotional signals shape perception in both brains and machines.

## Introduction

Emotion fundamentally shapes how we perceive and interpret the world around us. The brain has evolved sophisticated mechanisms to process the emotional information embedded in the sensory input. In vision, rather than proceeding as a purely bottom-up process, visual perception of emotion is dynamically modulated by internal emotional states and externally generated goals (Vuilleumier 2005; Pourtois et al. 2013). This flexibility enables humans to anticipate and prioritize emotionally salient information, disambiguate noisy inputs, and adaptively respond to complex visual scenes (Pessoa and Adolphs 2010; L. f. Barrett and Bar 2009). Anatomically, this operation is supported by recurrent connections between anterior emotion-modulating regions, such as the amygdala and ventromedial prefrontal cortex (vmPFC), and various hierarchical levels of the visual system (Cardinal et al. 2002; Amaral et al. 2003; Catani et al. 2003; Vuilleumier 2005; Pessoa 2009). Functionally, these bidirectional pathways underlie such distinct forms of emotion modulation as emotion expectation and emotion appraisal (Arnold 1960; C. A. Smith and Ellsworth 1985; C. A. Smith and Lazarus 1993; Sander et al. 2005; Cunningham and Brosch 2012; Moors et al. 2013; Yeo and Ong 2024). Impairments of these mechanisms are characteristic of many psychiatric disorders, including depression and schizophrenia. Thus, understanding recurrent processing in the visual-emotion circuit has both basic science and clinical significance.

Computational modeling has always played an active role in our pursuit to understand visual emotion processing. Recent advances in AI-inspired computational models are taking the field in a new and promising direction. However, the current deep learning models of emotion inference, though effective in static visual classification, rely on predominantly feedforward architectures (Krizhevsky et al. 2012; LeCun et al. 2015; Simonyan and Zisserman 2015; He et al. 2016; Kragel et al. 2019). They treat emotional perception as a direct mapping from image features to affective categories, overlooking the recursive, context- and goal-oriented dependent aspects of emotion perception (L. F. Barrett and Simmons 2015; Pei et al. 2024; Maniquet et al. 2024). As a result, they are not able to capture phenomena such as emotion anticipation and emotion regulation (Cunningham and Brosch 2012; R. Smith and Lane 2015). Bridging this gap calls for novel architectures that incorporate feedback and top-down mechanisms that enable context and internal state dynamics to better approximate the brain’s mechanisms for affective perception.

Recent NeuroAI work in the non-emotion domain has taken steps in this direction. For example, drawing inspiration from bidirectional cortico-cortical connections in the brain, Konkle and Alvarez (2023b) proposed a deep neural network architecture that incorporates long-range modulatory feedback connections to examine the influence of top-down cognitive steering on visual object recognition. Feedback pathways, along which top-down signals travel from high-level areas to early visual areas (Fişek et al. 2023; Gilbert and Li 2013), enable higher-level representations or external goals to dynamically influence lower-layer activations during recurrent inference (Lamme and Roelfsema 2000; Roelfsema and de Lange 2016). By integrating both internally generated and externally guided steering signals, architectures with feedback properties improve recognition performance for natural images, especially in ambiguous and target–distractor conditions, demonstrating the important role of feedback in solving the object recognition problem under environmental uncertainty. However, these types of models remain restricted to feedback within the visual hierarchy and do not address interactions between the visual and affective systems, which, as discussed earlier, are critical for visual emotion perception (Pessoa and Adolphs 2010; Sander et al. 2005; Pourtois et al. 2013).

In the present work, we attempted to address this problem by introducing EmoFB, a biologically inspired deep neural network model of human emotion perception. EmoFB consists of a visual system module and an affective system module, and the two modules interact bidirectionally. EmoFB supports two functionally distinct feedback routes: intrinsic feedback, arising from internal affective appraisals based on the model’s own perceptual representations, and external steering, which reflects broal contextual priors such as those derived from prior information or emotional anticipation. These higher-level signals are projected in a top-down fashion from the affective module, e.g., the amygdala–prefrontal system, back into the convolutional layers within the visual module recursively, forming a hierarchical cascade of modulatory signals that influence sensory processing. We evaluated EmoFB across multiple visual emotional perception tasks with varying levels of uncertainty and stimulus degradation, and assessed the underlying neural mechanisms using representational geometry. In addition, fMRI data recorded from participants viewing affective pictures were analyzed and compared with the deep neural network model to examine whether the introduction of feedback connections and bidirectional interactions helps to increase model-brain alignment.

## Results

### The EmoFB network model: Architecture, training, and performance

Prior deep neural network models of emotion perception contain only feedforward connections linking visual input to emotion categorization. In the biological brain, it is well-established that anterior emotion-sensitive brain areas both receive input from the visual system and send top-down signals to modulate visual processing (Vuilleumier 2005; L. f. Barrett and Bar 2009; Pessoa and Adolphs 2010). The EmoFB network, shown in Fig. 1A, builds on our previous feedforward Visual Cortex Amygdala (VCA) model (P. Liu et al. 2025), which demonstrated the effectiveness of coupling a visual system module with an affective system module. EmoFB extends this framework by connecting the two modules reciprocally with both feedforward and feedback connections. Specifically, in our model, the visual system module is based on a deep convolutional neural network (AlexNet) pretrained on recognizing natural images (Krizhevsky et al. 2012). It has been shown that the rich hierarchical representations of visual input in the AlexNet architecture parallel those of the human brain (Agrawal et al. 2020). For emotion processing and recognition, we modified the network by replacing its final object classification layer (1000 units) with a three-layered fully connected affective system module, which was then trained to recognize 8 discrete categories of emotion (Yang et al. 2023a).

**Fig. 1.**
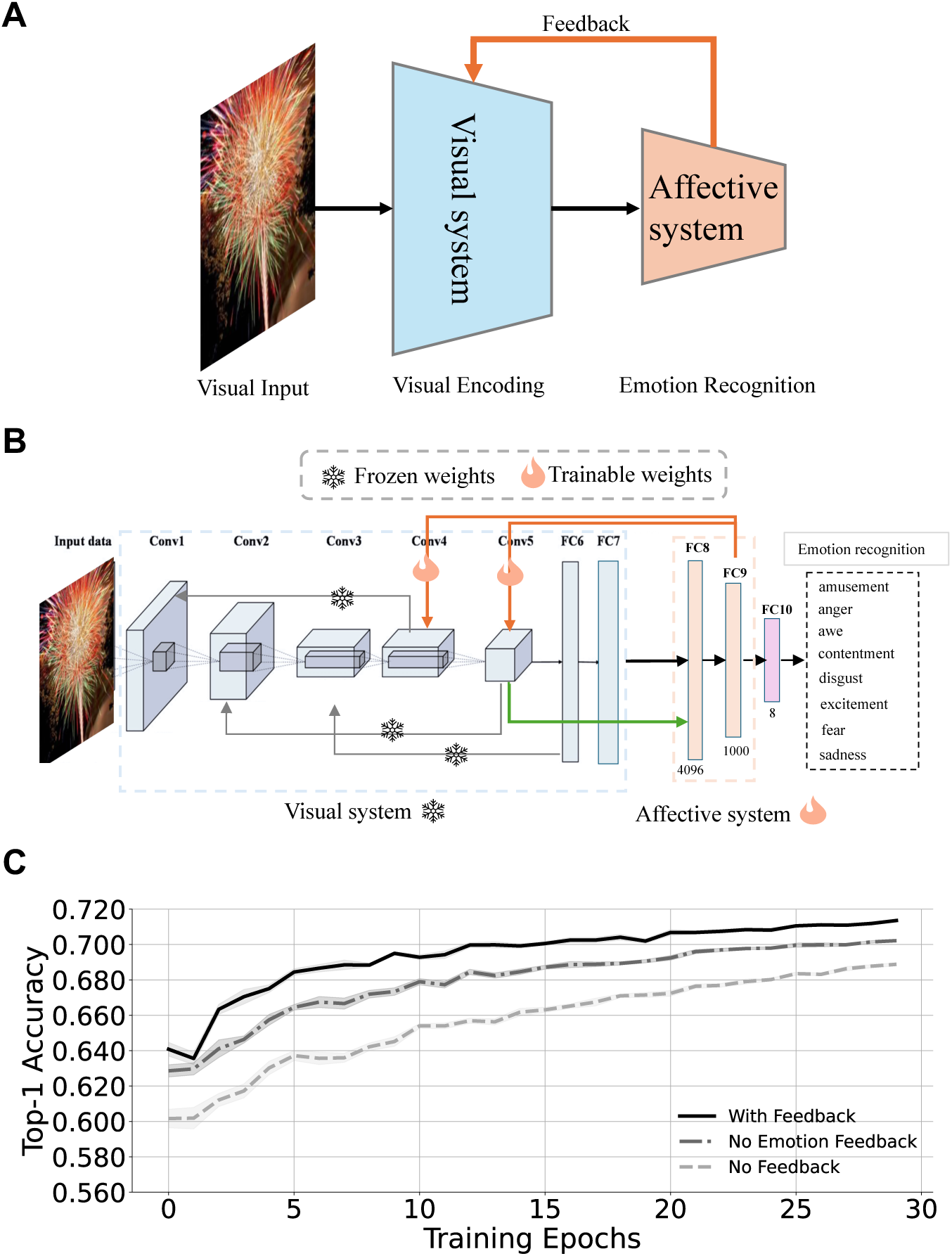
Architecture and training dynamics of the EmoFB network. **A.** The EmoFB model consists of two modules: a visual system module and an affective system module. The schematic illustrates that in addition to the conventional feedforward pathway, thee is a reentrant feedback pathway, allowing the modulation of the visual system by the affective system. **B.** Detailed connectivity within the EmoFB model. In addition to the layer-to-layer feedforward connection, feedback connections from FC9→Conv5 and Conv4→Conv2 are included, as well as an additional feedforward connection from Conv5→FC8. **C.** The model was trained to recognize the discrete emotion portrayed in the input image. Model performance as a function of learning: Top-1 accuracy over epochs for three model variants: full model with emotion-related feedback (black), no emotion-related feedback (gray solid), and no feedback (gray dashed). Models with full emotion feedback consistently outperform the others throughout training although the performance difference is quite modest. Shaded regions represent ±SEM (standard error of the mean) across 10 independently initialized EmoFB networks. FC: fully connected; Conv: convolutional.

As shown in Fig. 1B, in addition to hierarchically organized feedforward pathways, we added another feedforward connection (green line) from Conv5 to FC8 to transmit relatively less processed visual features into the affective system. This connection, after network training for visual object recognition, was fixed during emotion recognition training to preserve the integrity of bottom-up visual features while allowing us to isolate the effects of feedback. To integrate top-down modulation into the predominantly feedforward architecture, we added two types of feedback connections: (1) fixed connections within the visual system (gray arrows, e.g., Conv5→Conv2), whose weights were trained during visual object recognition and then held frozen during emotion recognition training, and (2) trainable connections from the affective system to the visual system (orange arrows, e.g., FC9→Conv5). The trainable feedback connections allow the affective system to learn to dynamically modulate visual representations based on high-level emotion recognition goals.

Training was carried out in two phases. The network was first trained on object recognition to establish stable visual representations. It was then trained on emotion recognition, during which the affective system module and its top-down feedback connections learned to modulate visual representations based on emotion classification demands. During the object recognition training phase, the visual system module, including hierarchically organized feedforward pathways and local feedback connections within the visual hierarchy, was optimized using a standard supervised learning objective. After this phase, all visual system module’s weights, including the Conv5→FC8 feedforward connection transmitting intermediate visual features to the affective system module, were frozen. Emotion recognition training was then performed in a supervised fashion to classify images into one of eight emotion categories: amusement, anger, awe, contentment, disgust, excitement, fear, and sadness (Yang et al. 2023a). During this phase, learning was restricted to the affective system module and the top-down feedback connections from the affective system module to the visual system module, allowing emotion-driven modulation of visual representations while preserving the integrity of bottom-up visual features. Two types of feedback are implemented in the EmoFB network. The first is intrinsic feedback, where the feedback signal is derived internally from the network itself—for example, from the activation of a deeper layer (e.g., FC9) in response to the current input stimulus (as shown in Fig. 1B). The second is external steering feedback, where the top-down modulation is guided by a category-level template signal (detailed in the next section). We first present results from intrinsic feedback below, and then examine the effects of external steering feedback.

As shown in Fig. 1C, networks equipped with top-down emotion-modulating feedback achieved consistently higher Top-1 accuracy throughout training compared to models with no feedback or with only object-recognition related feedback connections although the improvement is quite modest. The model with only object-recognition related feedback connections (gray solid line) performed better than the pure feedforward, no-feedback model (gray dashed line), indicating partial benefits from the local feedback within the visual system. Shaded regions denote ±SEM across 10 independently trained models (i.e., from 10 different random initializations), confirming the reliability of performance improvements driven by emotion-based feedback (see Method for more details).

### Top-down goal-oriented steering: Procedure and performance

As introduced above, the second type of feedback in EmoFB is external steering, where top-down modulation is guided by a template-like signal. For the task of recognizing whether a visual stimulus belongs to a specific emotion category, the steering signal is computed by averaging the FC9 activations across all images from the same emotion category in a held-out validation set (Fig. 2A). This averaged feature vector serves as a prototype representation of the target emotion, or a “template” (Duncan and Humphreys 1989; Desimone and Duncan 1995), and is injected into the model’s early layers to bias its visual processing.

**Fig. 2.**
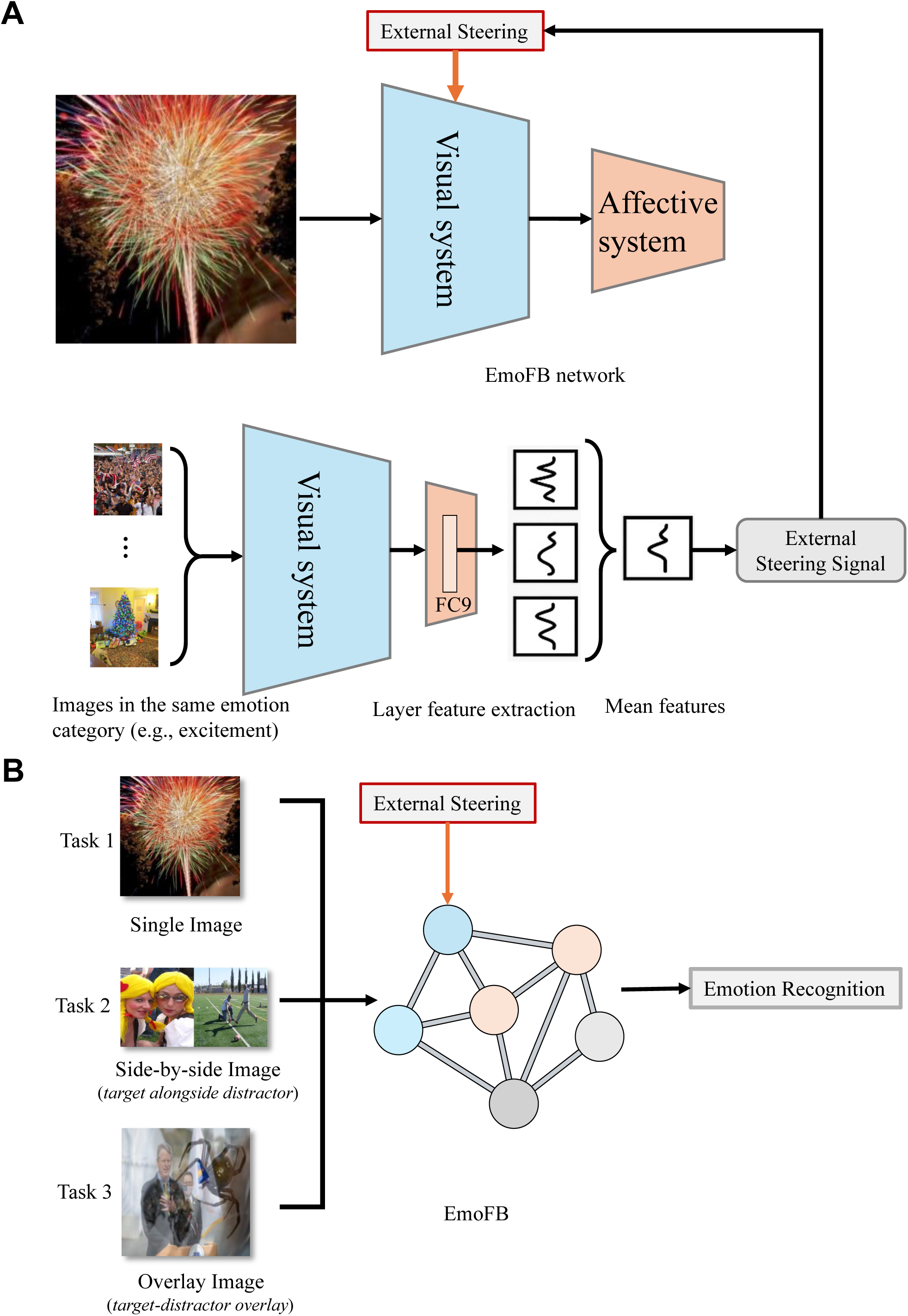
Model performance evaluation paradigm with external steering. **A**. Schematic of the EmoFB network with external steering. The external steering signal is constructed by averaging response patterns in layer FC9 (penultimate layer) evoked by images from the same emotion category as the visual input. This steering signal is fed back to the visual system module to modulate visual feature representations via top-down feedback. **B**. The network’s performance is tested on three input formats—single image (target only), side-by-side image (target + distractor), and overlay image (target-distractor superposition)—to evaluate emotion recognition under varying levels of visual ambiguity. Example images shown are from the EmoSet dataset(Yang et al. 2023b), a publicly available large-scale visual emotion dataset.

We tested how this external steering signal influences the network’s emotion recognition performance across three tasks of increasing visual complexity: (1) single image, where only the target image is shown; (2) side-by-side image, where the target is presented next to a distractor; and (3) overlay image, where the target and the distractor are fused into a single composite image (Fig. 2B). By applying external steering in each condition, we evaluate whether category-level top-down steering feedback enhances emotion recognition accuracy, even under strong distracting influence.

To evaluate the role of top-down modulation during recursive inference, the model performed up to five sequential feedforward-feedback passes (Fig. 3A), mimicking the brain’s recurrent computation between lower and higher level brain areas (Lamme and Roelfsema 2000; Gilbert and Li 2013). In the first pass, the network processes the visual input without top-down steering. In subsequent passes, feedback signals derived from the previous pass (intrinsic feedback) or from the template are applied to modulate target layers, progressively refining the network’s internal representational state. Each pass, therefore, builds on the prior one, allowing feedback to iteratively shape the representation before the final emotion recognition output.

**Fig. 3.**
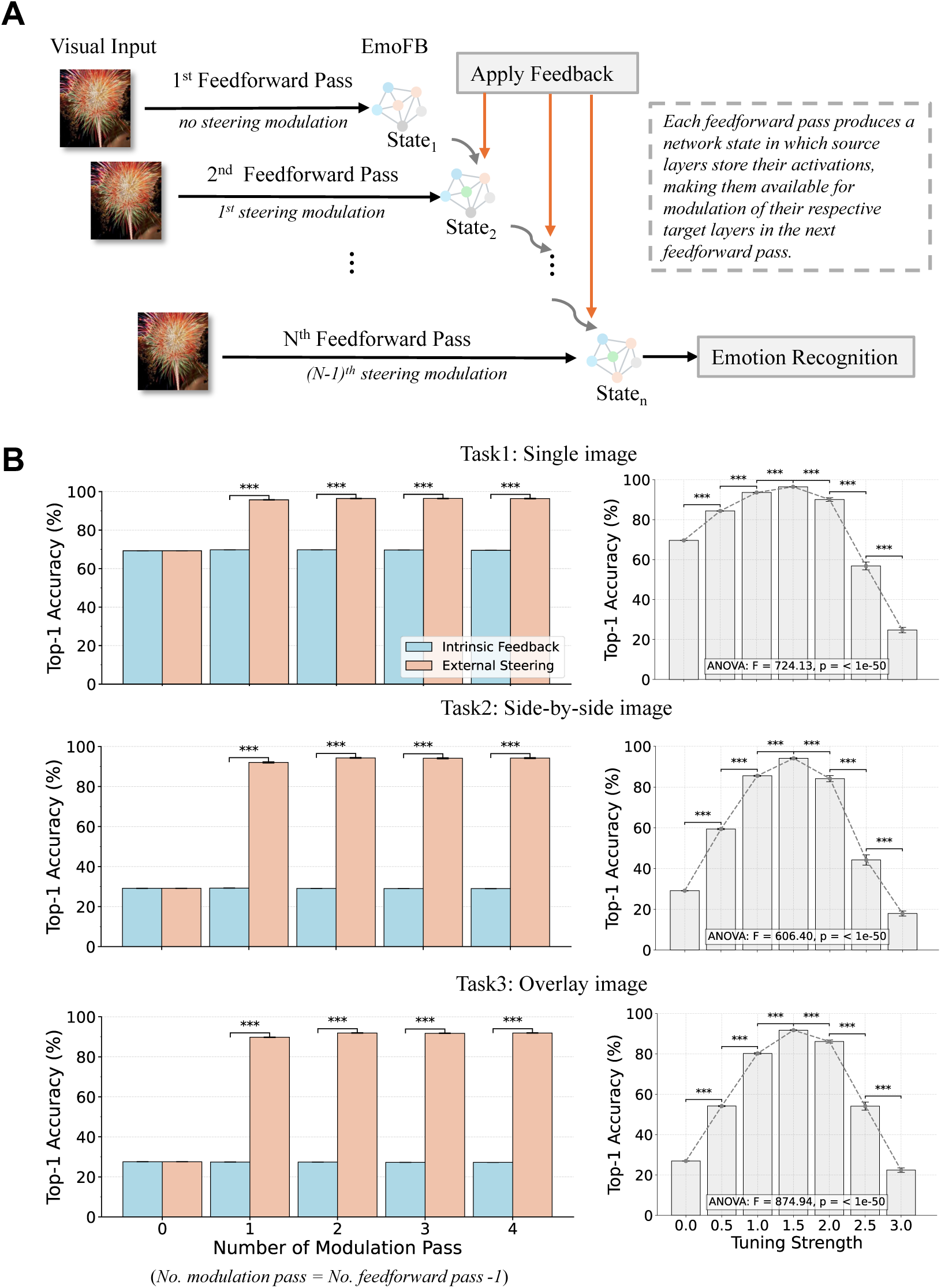
Model evaluation procedure and performance. **A.** Schematic of the behavioral testing algorithm. The EmoFB model performs multiple feedforward passes, with steering modulation (intrinsic or external) introduced after the first pass. Each feedforward pass generates a network state that modulates the next pass, enabling recurrent top-down influence. **B.** Top-1 emotion recognition accuracy across the three tasks. Left panels: Accuracy comparisons between intrinsic feedback and external steering across increasing numbers of modulation passes. Right panels: Accuracy as a function of steering tuning strength. Each bar reflects the mean ± standard deviation across 10 independently trained models initialized with random weights. Statistical significance was assessed using Welch’s t-test (two-sided, unequal variances) and ANOVA followed by post hoc tests. (***p < .001, **p < .01, *p < .05).

Across all three tasks, top-down steering consistently outperformed intrinsic feedback (Fig. 3B, left). In the most challenging experimental condition, in which the target image and the distractor image are superimposed, intrinsic feedback plateaued below 40% even after four modulation passes, whereas top-down steering drove performance above 80% after a single pass and over 90% at its peak performance. This highlights the effectiveness of top-down steering, which provides a category-prototype prior that helps disambiguate complex inputs. In addition, the magnitude of improvement scaled with task difficulty. For example, in the single-image task (low ambiguity), external steering improved accuracy by ∼20% over intrinsic feedback; in the side-by-side condition (moderate ambiguity), the improvement widened to over 65%; and in the overlay condition (high ambiguity), accuracy jumped from ∼27% under intrinsic feedback to ∼92% with external steering.

Interestingly, with external steering applied, recognition performance saturated after three feedforward passes (two modulation passes), with little or no gain from additional iterations. At this point, accuracy converged across tasks, 91% for side-by-side and 94% for overlay images, closely matching the 96% achieved in the single-image normal (ceiling-level) condition. This plateau suggests that the network rapidly settles into a stable representational state once feedback has reorganized the pattern based on early visual activations. A similar pattern is seen in biological vision, where (Lamme and Roelfsema 2000) noted that the feedforward sweep through the visual hierarchy is completed within ∼100 ms, after which recurrent feedback operates within a limited window to stabilize perception. Moreover, (Wyatte et al. 2012) showed that feedback strengthens degraded inputs until a stable state is reached, with further cycles offering little additional benefit. Thus, top-down goal-oriented steering functions as a rapid refinement mechanism, yielding most improvements in the first few iterations. These findings show that external steering feedback not only improves recognition when inputs are ambiguous but can also restore degraded perception in the distractors to the normal levels, paralleling the role of recurrent feedback in the brain in resolving uncertainty and stabilizing perception.

As discussed above, top-down steering based on category specific priors showed its greatest benefit under distracting conditions, a pattern that can be seen in biological vision. When sensory evidence is weak or uncertain, top-down signals from prefrontal and limbic regions shape perception by relying on prior expectations to resolve ambiguity (Summerfield and Egner 2009; Panichello and Buschman 2021). EmoFB captures the same principle. When input is clear, feedback adds only minor improvements, but when input is ambiguous, it becomes essential, adjusting internal representations toward the target or expected category, reducing the impact of distractors, and restoring recognition accuracy. In this way, the model offers a mechanistic account of how the brain relies on feedback most strongly when sensory evidence is unreliable, helping to stabilize perception in uncertain environments.

We further tested how top-down steering strength affected performance (Fig. 3B, right), mimicking how the brain regulates the gain of top-down signals to balance stability and flexibility (Grossberg 1980; Friston 2005). Accuracy improved as tuning strength increased from 0, reaching its best around ∼1.5, but declined when the strength was pushed further. This shows that top-down steering works best within an optimal range of feedback strength, where both too little or too much lead to suboptimal performance. This inverted-U pattern is similar to the Yerkes–Dodson law observed in biological systems (Yerkes and Dodson 1908), which showed that there is an optimal range of arousal that supports the best performance; very low or very high arousal both lead to degraded performance. Later studies extended this idea to cognitive control, showing that neuromodulators such as norepinephrine and dopamine are most effective at intermediate levels, with disrupted performance resulting when the levels are too low or too high (Grossberg 1980; Aston-Jones et al. 1999; Aston-Jones and Cohen 2005; Cools and D’Esposito 2011). At the systems level, (Reynolds and Heeger 2009) found that the effect of attention depends on how strongly neurons are already responding. Attention has the largest impact at medium stimulus contrasts, when responses are most flexible, but little effect once responses saturate at high contrast. This supports a general rule that top-down signals are most effective within a limited range. In line with these findings, our results show that feedback in EmoFB is most helpful at moderate tuning strength, while weaker or stronger signals reduce accuracy.

We reiterate that all results above are averaged over 10 independently initialized models, which enables significance testing using repeated-measures ANOVA, followed by post hoc comparisons. Together, the findings highlight that both the source (intrinsic vs. external) and the strength of top-down feedback are key determinants of emotion recognition. Notably, external steering allows the model to achieve near-ceiling level accuracy even under highly ambiguous conditions, paralleling the role of recurrent feedback in biological vision in resolving uncertainty and stabilizing perception.

### Neural mechanisms revealed by representational geometry

What are the neural mechanisms underlying the improved performance by top-down steering? To assess how top-down modulation shapes the internal structure of visual representations, we compared the model’s layer-wise representational similarity matrices (RSMs) with a theoretical RSM (TRSM) that encodes idealized category structure (Fig. 4A). In this theoretical RSM, images from the same emotion category are maximally similar (i.e., similarity=1), whereas images from different emotion categories are maximally dissimilar (i.e., similarity=0), resulting in a block-diagonal pattern. While this formulation does not capture graded similarities between different emotion categories, it provides a clear normative reference for evaluating the extent to which top-down modulation sharpens category separability. Pearson correlations between model-derived RSMs and the TRSM were computed across layers under both intrinsic feedback and external steering and evaluated separately for the three tasks (Fig. 4B).

**Fig. 4.**
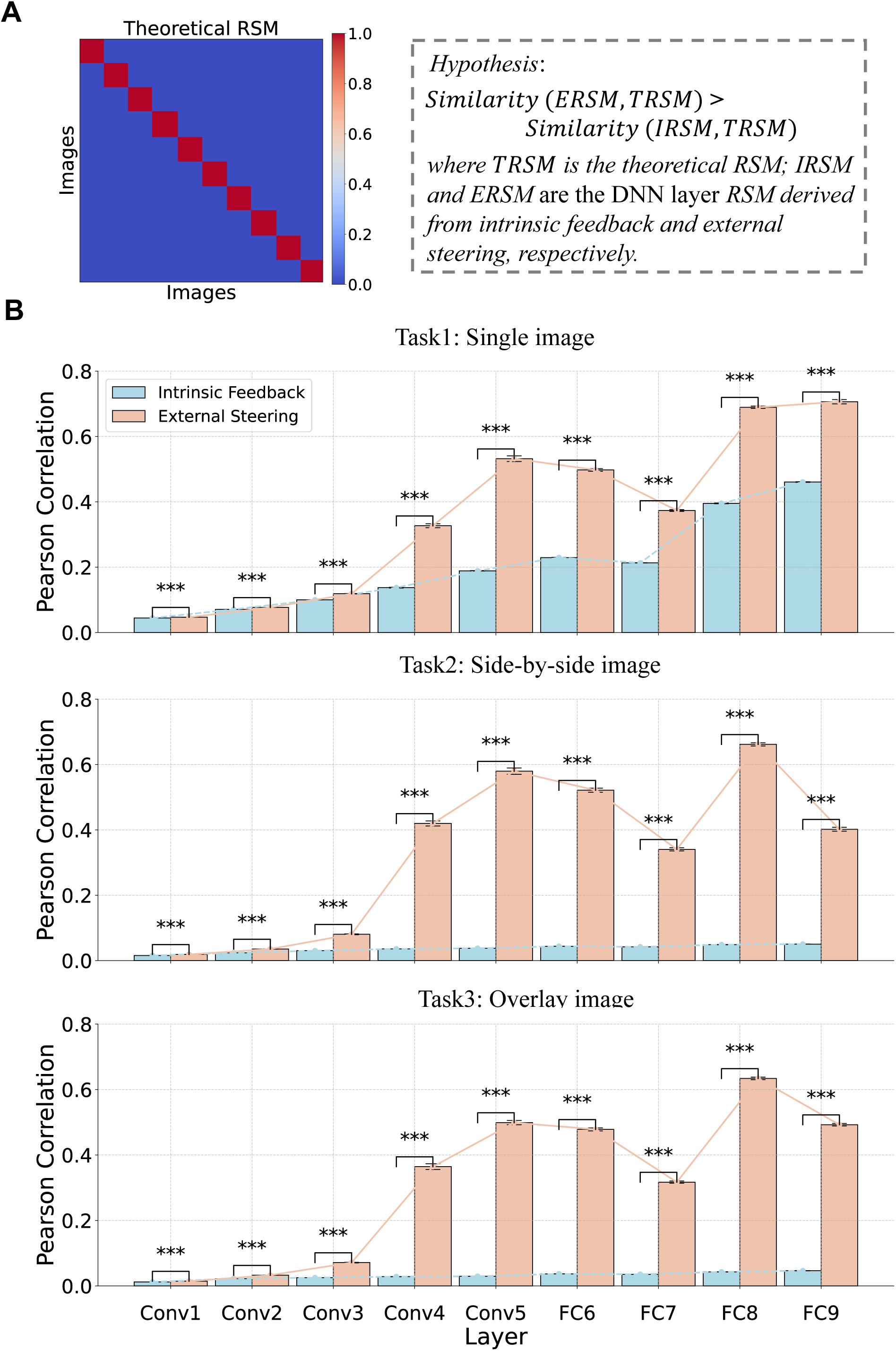
Neural representational geometry. **A.** Theoretical representational similarity matrix (TRSM) constructed based on categorical identity, where each image shares maximum similarity with others from the same emotion category (diagonal blocks). This matrix serves as a hypothesis-driven benchmark for evaluating model representations. **B.** Layer-wise representational similarity between the model’s RSMs and the theoretical RSM across three tasks: single image (top), side-by-side image (middle), and overlay image (bottom). Pearson correlation coefficients were computed between the theoretical RSM and the model-derived RSMs from each layer under intrinsic feedback (blue) and external steering (orange). Bars reflect the mean ± standard deviation across 10 independently trained models. External steering consistently enhances representational alignment with the theoretical structure, particularly in deeper layers (FC6–FC9) and under increased visual ambiguity. Statistical significance was assessed using Welch’s t-test (two-sided, unequal variances); asterisks indicate significance levels (***p < .001, **p < .01, *p < .05).

Across all conditions, external steering consistently enhanced representational alignment with the theoretical RSM, particularly in deeper layers and in tasks with greater visual ambiguity. In the single-image task, correlations under external steering increased steadily across layers, peaking at r = 0.69 (FC8) and r = 0.71 (FC9), compared to r = 0.40 and r = 0.46, respectively, under intrinsic feedback. Mid-layer gains were also substantial (e.g., Conv5: 0.53 vs. 0.19). In the more difficult side-by-side task, the representational correlation peaked at r = 0.66 (FC8) under external steering, while intrinsic feedback remained near zero (r ≈ 0.05) across all layers. Similarly, in the highly ambiguous overlay task, external steering yielded r = 0.63 (FC8) and r = 0.49 (FC9), while intrinsic feedback again failed to form meaningful category structure (r ≤ 0.05 throughout). These effects were statistically significant at nearly every layer (Welch’s t-test, ***p < .001), with 3–10× greater correlations under external steering relative to intrinsic feedback in the mid-to-late layers.

These results suggest that top-down steering reorganizes internal geometry by tightening within-category similarity and separating categories more clearly, thereby restoring meaningful structure even when sensory input is degraded and in the presence of distractors. Crucially, this reorganization mirrors the model performance results (Fig. 3). Just as external steering pushed recognition accuracy to near-ceiling (normal) levels under ambiguity, it also imposed a category-consistent structure on internal representations. In other words, the model performance gains emerge from representational restructuring, where top-down priors guide the network toward stable and semantically organized geometry despite noisy input.

### Model–brain alignment assessed via representational geometry

To evaluate whether EmoFB captures human-like neural representations, we performed representational similarity analysis (RSA) between model-derived and fMRI-derived similarity matrices (RSMs) using 60 affective images (20 pleasant, 20 neutral, 20 unpleasant). RSMs were computed before steering (after the first feedforward pass) and after steering (after the final feedforward pass), and then compared to RSMs from different brain regions of interest (ROIs). Following prior work on brain and deep neural networks correspondence (Yamins et al. 2014; Khaligh-Razavi and Kriegeskorte 2014; Cichy et al. 2016; Pham et al. 2023), early convolutional layers (Conv1–Conv2) corresponded to V1, mid-level layers (Conv3–Conv4) to V4, and higher fully connected layers (FC5–FC7) to Lateral Occipital Complex (LOC). The highest affect-related layers (FC8–FC9) were compared with the amygdala, consistent with recent work showing that artificial neural networks can model human amygdala responses to emotional stimuli (Jang and Kragel 2025). This mapping reflects the known gradient from low-level features in V1, to object-level coding in LOC, to affective evaluation in the amygdala (Amaral, Behniea, et al. 2003).

As shown in Fig. 5B, external steering increased model–brain similarity across most regions, with the largest gains observed in higher-level visual and affective areas. In the early visual cortex, similarity increased modestly. *V1–Conv1* rose from r = 0.19 to r = 0.21 and *V1–Conv2* from r = 0.23 to r = 0.26. In mid-level visual cortex, the improvements were more pronounced. *V4–Conv3* improved from r = 0.28 to r = 0.31 and *V4–Conv4* from r = 0.28 to r = 0.32. In higher-level regions, changes were subtler but still significant in some cases. *LOC–FC5* remained stable (r = 0.27 before and after steering), *LOC–FC6* increased from r = 0.35 to r = 0.37 (95% CI = [0.02, 0.01]), and *LOC–FC7* showed no change (r = 0.36 both before and after steering). The most notable effect was observed in the amygdala (FC8, FC9) comparison, where model–brain similarity improved from r = 0.22 to r = 0.27, representing a robust and statistically significant increase (95% CI = [0.06, 0.03], *p < .001).

**Fig. 5.**
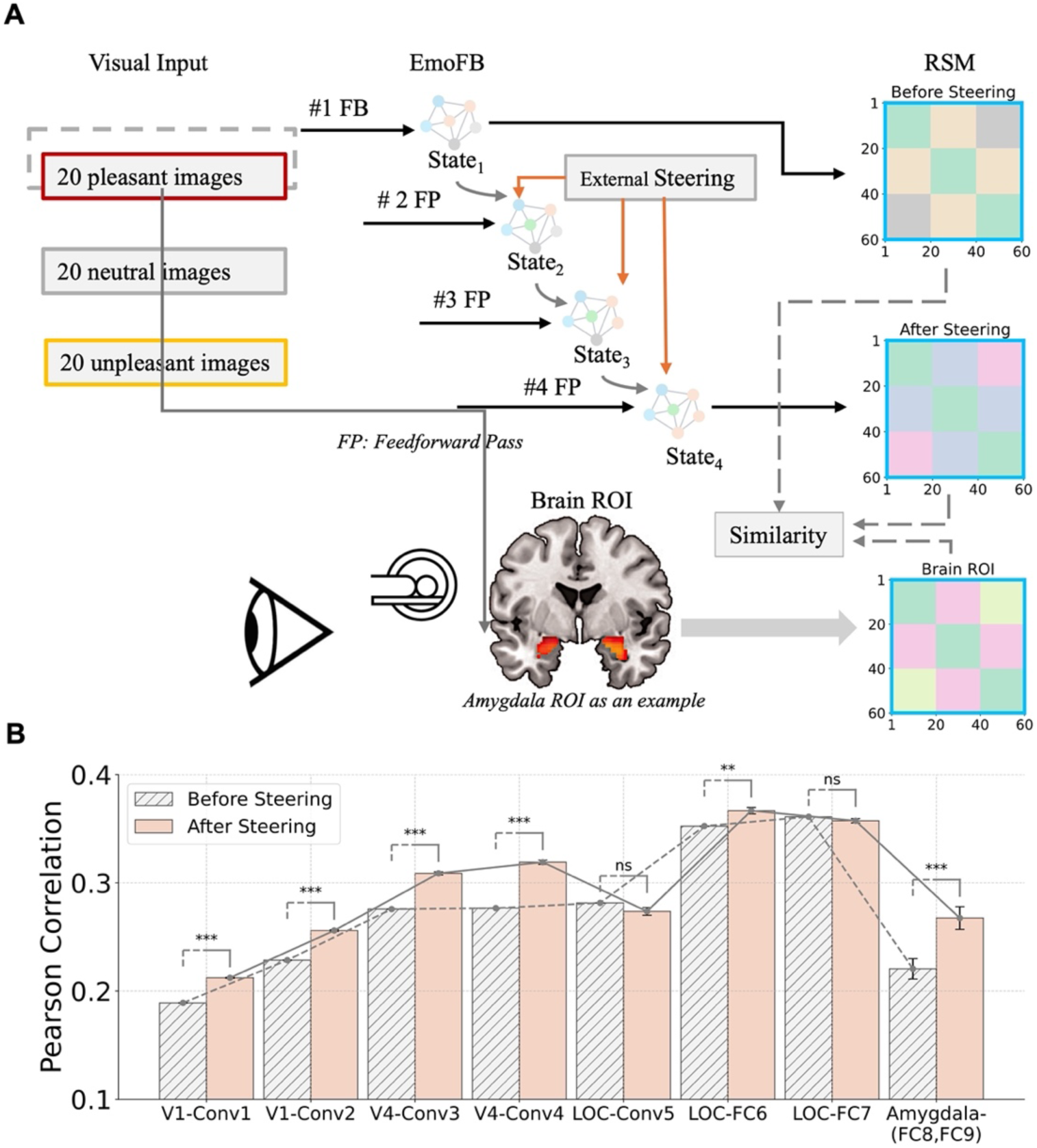
Comparing neural representations in the brain and EmoFB network. **A.** Schematic of the brain-model representational similarity analysis (RSA). Sixty affective images (20 pleasant, 20 neutral, 20 unpleasant) were presented to both the EmoFB model and human participants during fMRI scanning. The EmoFB model generated two representational similarity matrices (RSMs): one before steering (after the first feedforward pass), and one after steering (after the final feedforward pass). Corresponding brain RSMs were computed from fMRI activity in predefined regions of interest (ROIs), such as the amygdala. Representational similarity was quantified as the Pearson correlation between model-derived and brain-derived RSMs. **B.** Model–brain similarity (Pearson correlation) across layer–ROI pairs, where each x-axis label indicates the specific pairing between a brain ROI and a corresponding EmoFB layer (e.g., V1–Conv1 compares the V1 ROI with Conv1 layer representations; Amygdala–(FC8, FC9) compares the amygdala ROI with concatenated features from FC8 and FC9). Bars show the mean similarity before steering (hatched) and after steering (solid), averaged over 10 independently initialized models. External steering significantly enhanced brain-model alignment in most regions, especially in later stages of visual processing and the amygdala. Error bars denote standard deviation. Statistical significance was assessed using Welch’s t-test (***p < .001, **p < .01, *p < .05, ns = not significant).

These findings provide two insights. First, they show that external steering reorganizes internal representations in a way that brings the model closer to the brain’s representational geometry, from low-level visual encoding to higher-order affective perception. Second, they mirror biological evidence that emotion-related feedback alters sensory coding throughout the visual hierarchy. For example, emotional context modulates early visual cortex (Vuilleumier 2005), sharpens category distinctions in ventral temporal regions (Kensinger and Schacter 2006), and engages the amygdala to evaluate the emotional relevance of stimuli (Pessoa and Adolphs 2010). Thus, the improvement in model–brain alignment under external steering supports the view that contextual priors can reshape distributed representations in both artificial and biological systems, stabilizing perception and enhancing the salience of emotionally meaningful categories.

### Top-down steering in advance of stimulus input

Up to this point, top-down steering is triggered by bottom-up stimulus input. In many neuroscience experiments, an anticipatory state is established before the stimulus is shown. To examine the effect of top-down modulation of “brain state” on stimulus processing, we evaluated EmoFB when an external category-level signal was injected into the visual system before the first feedforward pass (Fig. 6A), referred to as pure external steering, conceptually resembling prestimulus neural activation by expectations, such as contextual- or category-specific priors conveyed from prefrontal or amygdala circuits to visual cortex (Bar et al. 2006; Summerfield and Egner 2009; Kok et al. 2017). In other words, unlike the recurrent computational setup studied earlier (Figs. 3–5), where steering follows the first pass and influences the network over multiple iterations, this condition tests whether a one-time provision of prior knowledge is sufficient to improve performance and reorganize internal representations.

**Fig. 6.**
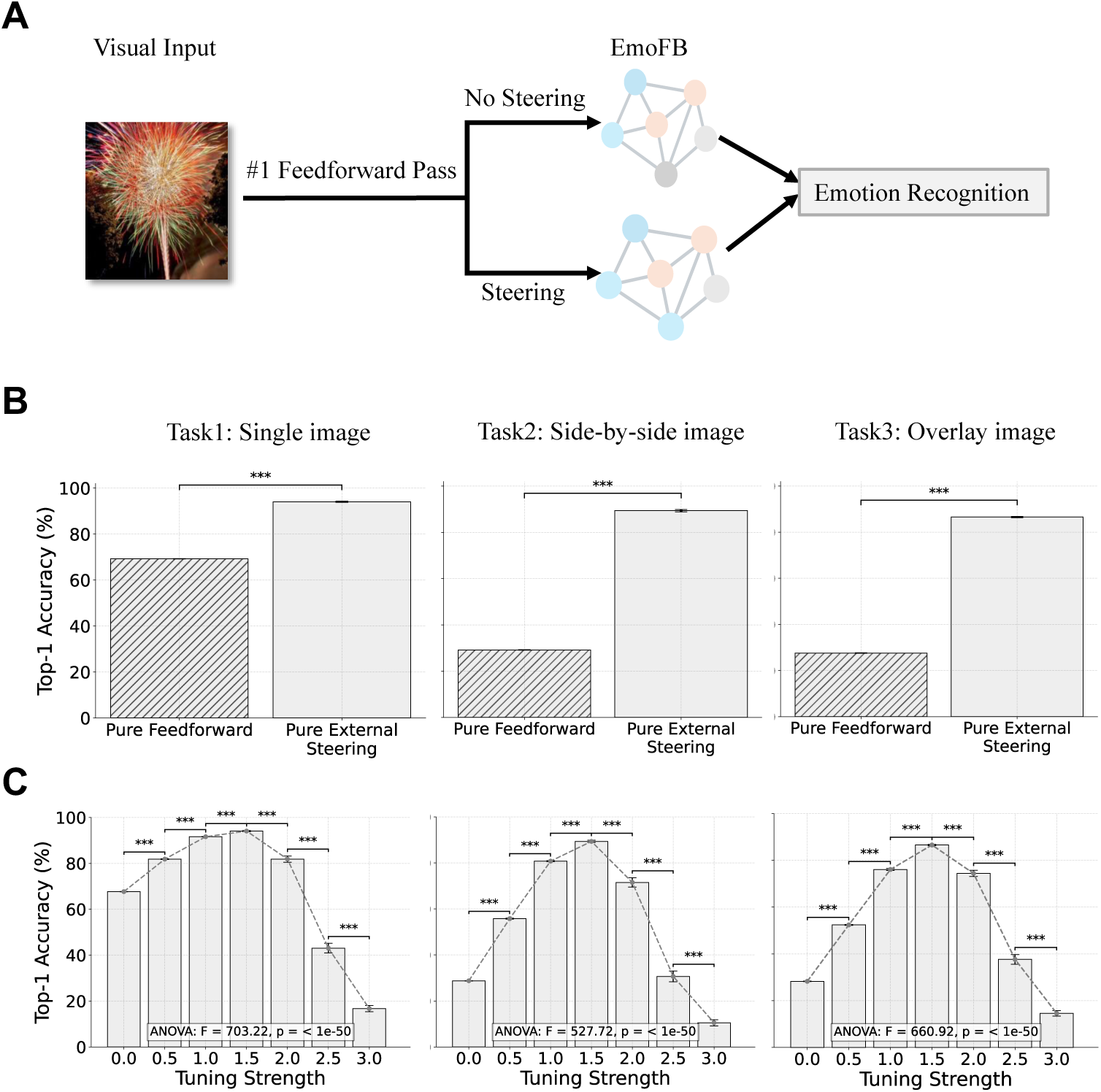
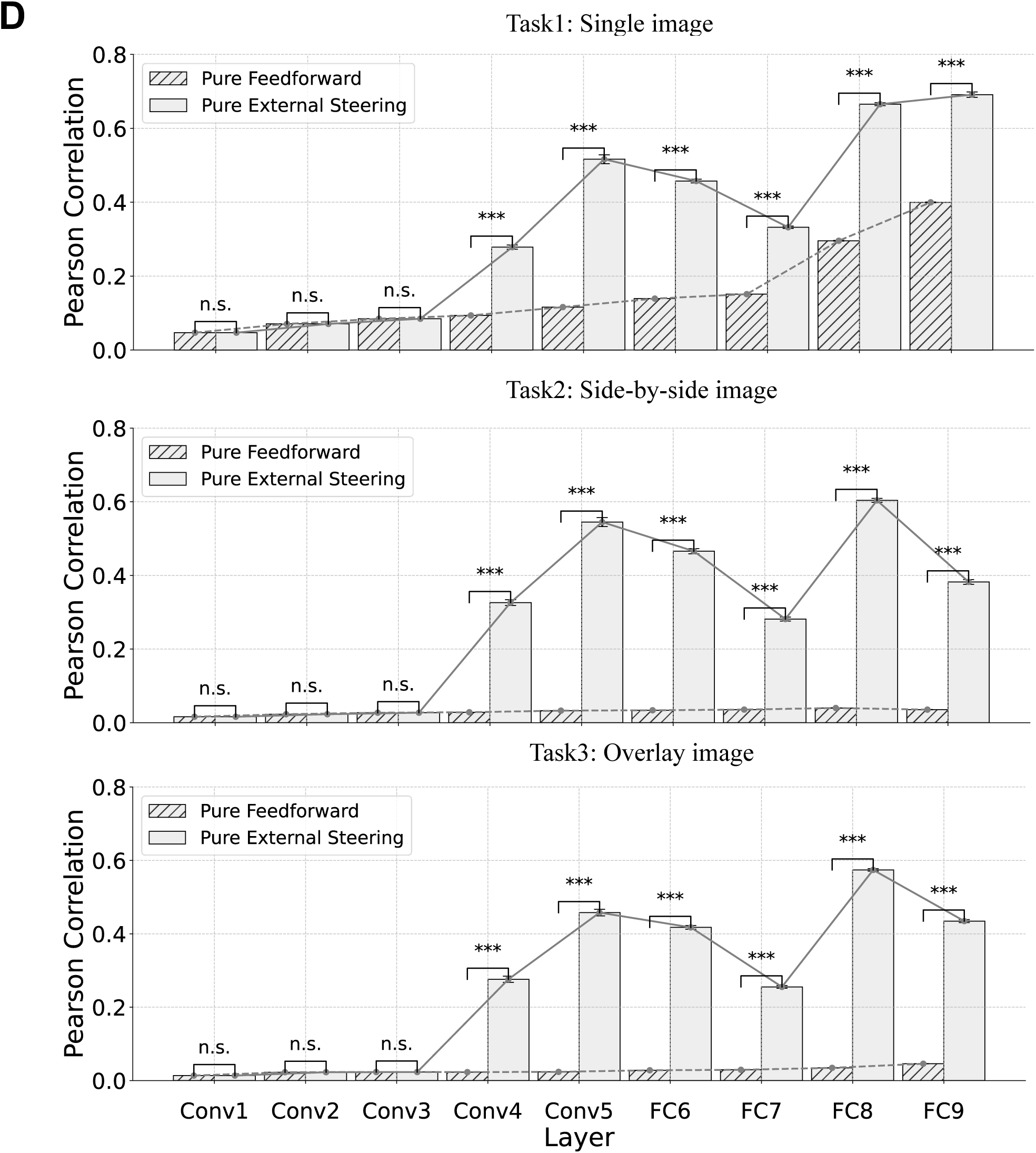
Pure top-down steering without recurrent modulation. **A.** Schematic of the pure top-down steering condition. Unlike the recurrent version used in previous figures, where steering is applied after the first feedforward pass, here, the external steering signal is injected before the first pass. Target layers are pre-activated by the steering signal, and the network performs only a single feedforward pass. No recurrent modulation is used. **B.** Top-1 emotion recognition accuracy across three tasks, single image (left), side-by-side image (middle), and overlay image (right), comparing pure feedforward (no steering) and pure external steering (steering applied only once at input). **C.** Accuracy as a function of tuning strength under pure top-down steering. **D.** Layer-wise representational similarity (Pearson correlation) between model RSMs and the theoretical RSM under pure feedforward vs. pure top-down external steering. Here, FC8 and FC9 are shown separately to reveal layer-specific effects of steering within the affective system module; in Fig. 5B, they were grouped by their shared brain ROI (amygdala) for model–brain comparison. Steering improves category structure in deeper layers, especially for more ambiguous tasks. Error bars denote the standard error of the mean across 10 model initializations. Asterisks mark statistical significance (Welch’s t-test: ***p < .001, **p < .01, *p < .05, ns = not significant).

Behaviorally, pure external steering produced significant gains in Top-1 accuracy across all three tasks compared to the pure feedforward condition (Fig. 6B). In the single-image task, accuracy increased from 69% to 92%. In the side-by-side condition, performance rose from 29% to 89%, and in the overlay condition, from 28% to 87%, roughly tripling of accuracy in the two ambiguous conditions the absence of recurrent processing. We next examined the effect of tuning strength on performance under pure external steering (Fig. 6C). Across all tasks, Top-1 accuracy followed an inverted-U profile, peaking at a tuning strength of 1.5. For example, in the overlay task, accuracy peaked at 87% at a strength of 1.5, then declined slightly at higher strengths. This pattern mirrors the behavior seen under recurrent steering, suggesting that optimal integration of priors is tunable even in a single-pass context.

Finally, we assessed how pure external steering influenced internal representational structure by comparing model RSMs with the theoretical category-based RSM (Fig. 6D). As in previous analyses, steering enhanced representational similarity in deeper layers, especially under ambiguous conditions. In the overlay task, Pearson correlation in FC8 rose from r = 0.03 to 0.57, and in FC9 from r = 0.05 to 0.43. In the side-by-side task, FC8 improved from r = 0.04 to 0.60, and FC9 from r = 0.04 to 0.38. Even in the single-image task, steering enhanced FC9 similarity from r = 0.40 to 0.69. All improvements were statistically significant (Welch’s t-test, ***p < .001).

Taken together, these findings demonstrate that anticipatory modulation of visual cortex implemented by top-down steering can markedly enhance emotion recognition and reinforce category-specific clustering in representational geometry, even without recurrent processing. This parallels behavioral priming in humans, where expectations pre-activate task-relevant representations, which in turn accelerate recognition and sharpen category structure.

### Effect of top-down steering on false positive rates

One drawback of top-down steering is that it increases false positive rates. We assessed how pure top-down external steering (i.e., steering applied before the first feedforward pass, as in Fig. 6) influenced recognition in target-absent trials, where the input images did not contain the steered emotion category (Fig. 7A). As shown in Fig. 7B, steering substantially increased false positive responses across all three task conditions. At tuning strength = 0, false positive rates were low, but they rose sharply with increasing tuning strength, peaking at intermediate values of 1.5 before declining slightly at stronger levels of steering. This inverted-U pattern indicates that steering progressively biases the model toward reporting the steered emotion, even when it is absent. The magnitude of this bias varied with task difficulty. False positives were most pronounced in the side-by-side and overlay conditions, reaching peak levels above 60–70%, consistent with the greater perceptual ambiguity in these tasks. In contrast, the single-image condition showed only a modest increase, with peak false positives remaining below ∼50%, an intriguing result given that, in the target-present case, steering yielded the strongest performance gains on ambiguous composite conditions (overlay and side-by-side) compared to single-image conditions (Fig. 3B, Fig. 6B). This asymmetry suggests that the effect of top-down steering scales with perceptual ambiguity. In all three conditions, steering substantially improved target-present recognition, but the improvement was greatest when competing emotional inputs increased perceptual uncertainty. Correspondingly, false positive rates in target-absent trials were also highest in these ambiguous conditions (Fig. 7B), indicating that the same ambiguity that allows steering to enhance detection also makes the model more susceptible to misattributing the steered emotion when it is absent. This pattern parallels human perception, where top-down expectations enhance detection under clear conditions but, in the presence of ambiguity, can lead to misperceptions, such as mistaking a vague shadow or blurred spot for a threatening stimulus like a spider or snake (Öhman 2005; Sterzer et al. 2008). Ambiguity amplifies the influence of priors, making false alarms more likely when multiple interpretations are possible.

**Fig. 7.**
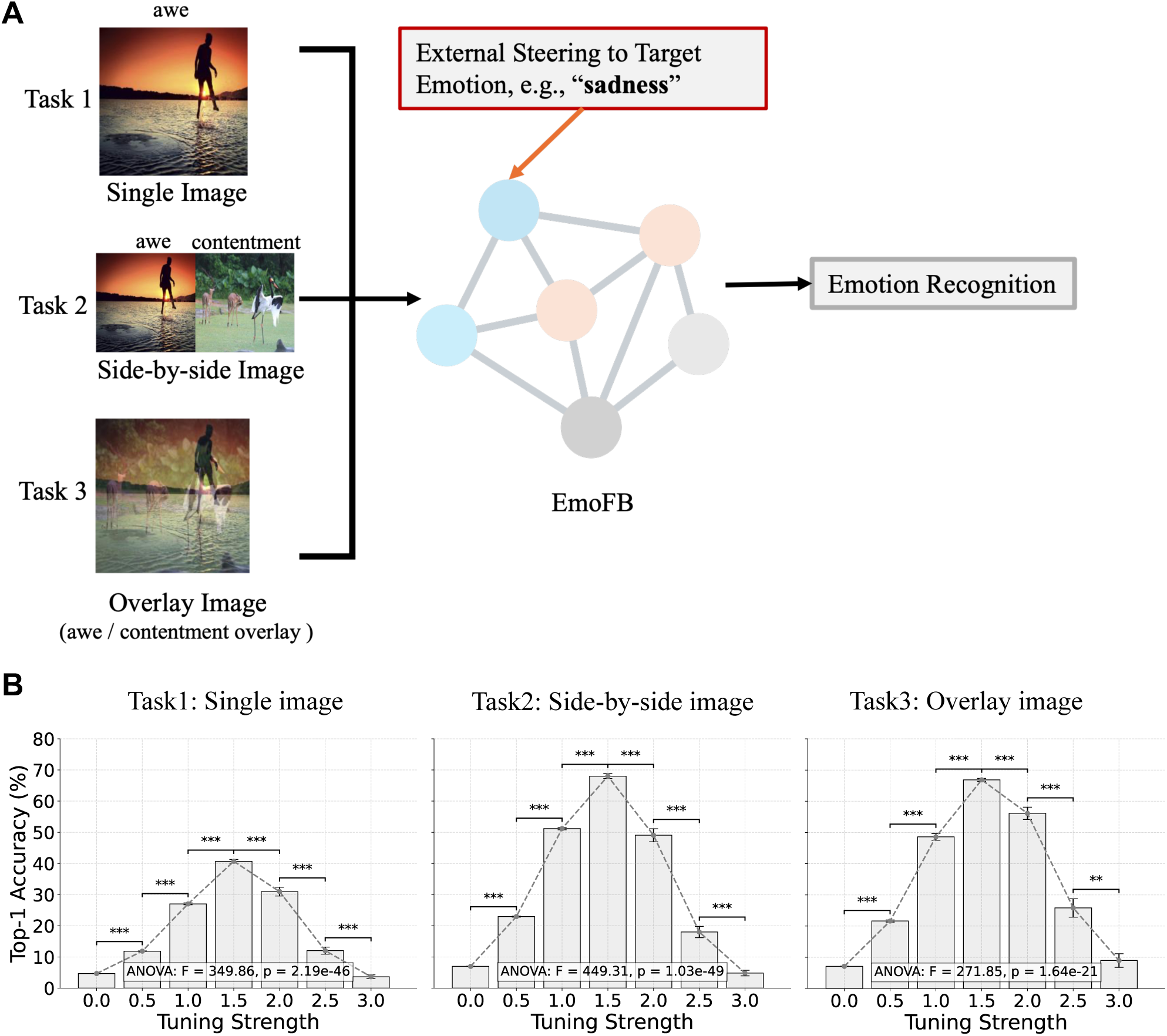
Pure external steering performance as a function of tuning strength in target-absent trials. **A.** Schematic representation of steering a target emotion category (e.g., sadness) in input images that do not initially exhibit the target emotion. **B.** Top-1 accuracy plotted as a function of tuning strength across three image display conditions: single, side-by-side, and overlay.

Thus, while external steering effectively enhances recognition when the target emotion is present, it also heightens the risk of false alarms in its absence, particularly under conditions of competing emotional content. Together, these findings highlight a fundamental trade-off of top-down steering. It increases sensitivity to target emotions but reduces specificity, with the greatest vulnerability to false positives emerging in ambiguous contexts. This trade-off provides a computational parallel to human emotion perception (Moratti and Keil 2009; Bradley et al. 2012; Gruss and Keil 2019), where expectation-driven feedback can sharpen recognition of relevant signals but also increase susceptibility to false alarms under uncertainty.

## Discussion

In this study, we introduced EmoFB, a deep neural network model designed to examine how top-down feedback mechanisms shape visual emotion recognition. The model integrates a feedforward visual system module with an affective system module through both bottom-up and top-down pathways, allowing high-level emotional representations to modulate early visual processing. This cross-system design, inspired by anatomical pathways linking the anterio emotion-modulating structures such as the amygdala and prefrontal cortex to visual cortex (Amaral, Behniea, et al. 2003; Pessoa and Adolphs 2010; Barbas 2015), extends prior models by embedding an explicit affective–visual loop that captures the dynamic interplay between appraisal and perception (Gilbert and Li 2013; Lamme and Roelfsema 2000; Roelfsema and de Lange 2016). Across tasks with varying levels of ambiguity, feedback improved emotion recognition, with externally guided steering providing the largest benefits. It greatly boosted accuracy, sharpened category structure in the model’s internal representations, and made these representations more consistent with fMRI responses in both visual and affective brain regions. Concurrent with these positives we also note the negative effect of external steering in terms of increased false positives.

### Emotion shapes perception

Emotion recognition is not just a feedforward process. In real-world contexts, emotion perception, which depends on the recognition of objects, faces, and scenes, often requires resolving ambiguity and integrating prior knowledge, expectations, and internal states (Vuilleumier 2005). Top-down feedback is central to this process; rather than passively encoding sensory data, the brain continuously reinterprets visual input in light of prior knowledge about its emotional relevance (Pourtois et al. 2013). EmoFB demonstrates this principle computationally. We show that feedback signals, constructed from emotion-category prototypes, reorganize intermediate visual features, improving recognition accuracy and restoring category structure even under ambiguous input. This mirrors neuroscience evidence that the amygdala and medial prefrontal cortex project back to visual areas, biasing processing toward emotionally salient information (Amaral, Behniea, et al. 2003; Vuilleumier and Driver 2007). Functionally, such feedback enhances detection of threat- or reward-related cues under uncertainty, tuning perception toward motivationally relevant features (Öhman 2005; Pessoa and Adolphs 2010; Pourtois et al. 2013). By integrating these behavioral and representational findings with neurobiological evidence, EmoFB highlights how emotional feedback can sharpen perception under ambiguity. The model thus provides a computational framework for testing hypotheses about how emotional states reshape sensory coding in the brain.

### Multiple forms of top-down control

EmoFB incorporates two complementary forms of top-down modulation: intrinsic feedback and external steering. Intrinsic feedback is appraisal-like and stimulus-driven, using activations within the affective module, derived directly from the current input, to modulate early visual layers. External steering, by contrast, is expectation-based, injecting category-prototype signals into early layers as state-dependent priors (Vuilleumier 2005; Pourtois et al. 2013). Our results show that both mechanisms improve performance. Whereas intrinsic feedback yields modest gains, external steering produces substantial accuracy improvements, especially under ambiguous conditions. This difference arises because intrinsic feedback is limited to the information in the current stimulus. Such signals resemble cognitive appraisal in the brain, where affective interpretations of incoming stimuli are formed and updated (Sander et al. 2005; Cunningham and Brosch 2012; Moors et al. 2013), but without the influence of learned priors or contextual cues. As a result, the intrinsic signal carries internal noise and is insufficient to drive great improvements in recognition when inputs are degraded or conflicting. External steering, by contrast, provides a clear and structured prior. Importantly, external steering can be applied in advance of stimulus input (referred to as pure external steering), paralleling anticipatory modulation in the brain, where task demands, contextual cues, and past experience bias perception toward anticipated emotional categories (Summerfield and Egner 2009; W. Li and Keil 2023). Neuroimaging and behavioral studies further show that emotional cues capture attention automatically but have much stronger and more reliable effects when they are directly relevant to current goals, reflecting interactions between prefrontal control signals and amygdala-based affective signals (Vuilleumier 2005; Pourtois et al. 2013). By injecting prototype-based signals, whether bottom-triggered or top-down applied in the absence of stimulu inpus, EmoFB reshapes internal representations and imposes categorical structure on early visual processing, enabling the network to resolve ambiguity and align perception with task-relevant emotional states.

### Model-brain representational alignment

Top-down feedback not only improved task performance but also increased the similarity between EmoFB’s internal representations and human brain activity. Using RSA, we found that external steering strengthened correlations between model and brain RSMs, particularly in V1, V2, LOC, and the amygdala. These results parallel the role of emotional top-down feedback in the brain. Emotional stimuli engage the amygdala, whose outputs influence activity in visual cortical areas, amplifying processing of emotionally salient information. fMRI and ERP studies report stronger responses to emotional versus neutral stimuli in early visual cortex (Lang et al. 1998; Pourtois et al. 2004; Padmala and Pessoa 2008; Y. Liu et al. 2011) and in higher-order regions such as fusiform cortex (Morris et al. 1998; Vuilleumier and Schwartz 2001; Sabatinelli et al. 2005). Anatomical and imaging studies provide a mechanistic basis for these effects, revealing bidirectional pathways linking the amygdala and prefrontal cortex with both early and mid-level visual areas (Amaral, Behniea, et al. 2003; Catani et al. 2003). Lesion studies further demonstrate causality: amygdala damage abolishes the visual enhancement normally observed for emotional stimuli (Vuilleumier et al. 2004).

The enhanced brain–model alignment observed in EmoFB suggests that its steering signals mimic this biological feedback mechanism. Notably, the effect extended from early visual regions to higher ventral areas, consistent with the view that affective feedback cascades through the visual hierarchy, shaping perception from its earliest stages (Amaral, Behniea, et al. 2003; Vuilleumier and Driver 2007). This mirrors our network architecture, where top-down pathways from the affective module modulate higher layers (e.g., Conv5) and recursively influence earlier ones (e.g., Conv2), echoing amygdala–visual feedback loops described in neuroanatomical studies (Pessoa and Adolphs 2010). Electrophysiological evidence supports this interpretation, showing that emotional stimuli boost very early visual responses, such as the first wave of activity in primary visual cortex and slightly later signals in nearby extrastriate areas, consistent with attentional gain-control mechanisms (Pourtois et al. 2004; Rotshtein et al. 2010; Y. Liu et al. 2011).

### False alarms in brains and machines

Although emotional top-down steering can substantially enhance emotion recognition, it can also increase false positives. In these cases, the steering signal biases the model to detect the target emotion even when it is not present in the stimulus. Our target-absent analysis (Fig. 7) showed that false alarms were not uniform across tasks; they were much higher in the side-by-side and overlay conditions than in the single-image condition. This pattern suggests that stimulus ambiguity amplifies the influence of top-down expectations, when multiple emotions compete within a scene, steering is more likely to bias recognition toward the cued category, even in its absence.

A similar phenomenon is well documented in the brain, where emotional states can bias perception and, through feedback, override bottom-up sensory input. For example, expectations about threat can shift the interpretation of ambiguous stimuli toward fearful meanings even in the absence of actual threat cues (Phelps 2006). Electrophysiological evidence further shows that emotional states amplify early visual responses, even when stimuli are neutral or degraded (Pourtois et al. 2004; Rotshtein et al. 2010). This pattern is consistent with an “expectation overweighting” mechanism, whereby affective priors exert disproportionate influence on perception, biasing ambiguous input toward anticipated emotional categories (Phelps 2006; Vuilleumier 2005; Pessoa and Adolphs 2010). In anxiety and post-traumatic stress disorder (PTSD), heightened expectations of threat amplify false alarms, leading individuals to misperceive neutral or ambiguous cues as threatening (Bishop 2007; Fani et al. 2012). Such biases illustrate how mechanisms that are adaptive in uncertain or threatening environments, where missing an emotional cue carries a greater cost than mistakenly perceiving one, can become maladaptive when feedback is chronically overweighted.

### Comparisons with related work

Most existing deep learning models for emotion recognition rely on feedforward architectures, treating emotional categories as static labels (Mollahosseini et al. 2019; S. Li and Deng 2022). Few attempt to capture how emotional interpretation dynamically reshapes perception. Even models that incorporate recurrence or attention generally lack explicit affective-to-visual feedback (Kollias and Zafeiriou 2020; Zhang et al. 2015). When feedback is included, it typically remains confined within a single processing system rather than crossing between affective and perceptual systems.

EmoFB is a biologically inspired model that explicitly incorporates top-down feedback between affective and visual systems. While informed by prior vision-based feedback models (Konkle and Alvarez 2023b), it advances them by introducing a dedicated affective system module and redirecting feedback from a purely visual–to-visual pathway to a visual–to–affective–to–visual pathway. In this way, EmoFB was designed to simulate the emotional attentional control mechanisms thought to operate in human perception. By introducing cross-system feedback from an affective module into a visual module, EmoFB implements a biologically inspired mechanism of top-down modulation. This design yields an interpretable architecture with emotion-specific feedback that goes beyond performance optimization, offering a principled framework for investigating how emotional states shape perception. EmoFB integrates prior work on visual feedback with a neurobiologically grounded account of emotion–perception integration.

### Summary

This study introduces EmoFB, a biologically inspired neural network that integrates an affective system with a visual processing system through top-down feedback. By allowing emotional representations to modulate visual layers, EmoFB captures the reciprocal influence between perception and emotion. The model implements two complementary feedback mechanisms: (1) intrinsic feedback, stimulus-driven and appraisal-like feedback; (2) external steering, bottom-up triggered and expectation-based feedback. Both improve recognition under ambiguity, with external steering providing the strongest gains. Representational analyses show that steering sharpens category structure and increases alignment with human brain activity, particularly in the early visual cortex, LOC, and the amygdala, underscoring the model’s biological plausibility. EmoFB thus provides a framework for testing how emotion-based feedback shapes perception across hierarchical layers, how internally versus externally guided cues influence recognition, and how feedback alters brain–model alignment, questions directly testable with fMRI, EEG, or behavioral paradigms.

## Materials and Methods

### Image datasets for model training and testing

Training and testing of the EmoFB network utilized two primary image datasets: ImageNet (Deng et al. 2009) and EmoSet (Yang et al. 2023b). The ImageNet dataset, a widely used dataset in computer vision and deep learning, was used to pretrain the visual system module for object recognition, enabling it to extract rich, general-purpose visual features from natural scenes. The EmoSet dataset, a more recently published dataset, was developed for emotion recognition. It contained images from diverse visual sources such as social media and artistic platforms (Yang et al. 2023b). EmoSet comprises over 3.3 million images, including 118,102 labeled images, each annotated with one of eight emotion categories: amusement, anger, awe, contentment, disgust, excitement, fear, and sadness. In this study, the labeled EmoSet data were divided into three sets: 80% training, 5% validation, and 15% test. Accordingly, we used 90,664 images for training, 7,998 for validation, and 19,440 for testing (see Supplementary Fig. S1A). To mitigate class imbalance during training, we applied class weighting based on the inverse frequency of each emotion category in the training set (Supplementary Fig. S1B).

To systematically evaluate model performance under different visual tasks, we constructed a composite test set of triplets from the EmoSet test images. Each triplet includes three presentation formats: original, overlay, and side-by-side (see Fig. 7A). The number of images per format was approximately balanced across emotion categories to ensure fair comparisons across conditions. Roughly 1,200 images were included in each format, forming the basis for all top-down feedback–related model testing and theoretical RSA analysis.

All input images were resized to 224×224 pixels. During training, we applied random resized cropping, random horizontal flipping, and ImageNet-style normalization (mean = [0.485, 0.456, 0.406]; std = [0.229, 0.224, 0.225], scaled by 255). For the test, images were center-cropped to the same resolution and normalized identically. Pixel values were cast to float16 for memory efficiency. Since image files are typically stored in Height × Width × Channel (HWC) format, all images were converted to Channel × Height × Width (CHW) format before being passed into the network, as required by PyTorch convolutional layers.

We additionally evaluated EmoFB using the International Affective Picture System (IAPS) (Bradley and Lang 2007), a standard stimulus set in affective neuroscience with normative ratings along continuous valence and arousal dimensions. IAPS images were not used for training or validation. They were reserved exclusively for post-training analyses to assess whether affective representations learned from EmoSet generalized to a canonical neuroscience dataset.

All IAPS images were preprocessed identically to EmoSet test images (224×224 resolution, ImageNet normalization, CHW format) and passed through the trained network in inference mode. Internal representations were extracted from selected layers and analyzed using RSA.

### Deep neural networks

The proposed EmoFB network consists of two modules: a visual system module and an affective system module (Fig. 1A–B). The visual system module was adapted from the study (Konkle and Alvarez 2023b), which was based upon AlexNet, but with the addition of feedback connections from the final fully-connected layer to earlier fully-connected and convolutional layers. The visual system was trained on ImageNet for object recognition. For our implementation, we retrained this module for 300 epochs on ImageNet using four NVIDIA A100 GPUs. Training was conducted with a batch size of 1024, an initial learning rate of 0.1 decayed by a factor of 0.1 every 50 epochs, and cross-entropy loss as the optimization objective.

For emotion recognition, we replaced the last fully connected layer for object recognition with the affective system module, which includes 3 fully connected layers. Feedbacks come from the second-to-the-last layer of the affective system module and connects to Conv5 and Conv4 layers of the visual module. The rationale for choosing the second-to-the-last layer instead of the last layer of the affective system module as the source of the top-down feedback signal is that this layer has richer emotion-related semantic features compared to the last layer, which has only 8 units corresponding to 8 discrete emotions (Yang et al. 2023a): amusement, anger, awe, contentment, disgust, excitement, fear, sadness.

To complete the EmoFB network model, we added one additional feed-forward connection from Conv5 of the visual system to the first layer (FC8) of the affective system. The information from this connection is integrated (concatenated) with the feed-forward connection from FC7 in the visual system. In this way, the affective system receives not only the high-level visual semantic features but also the less-processed features, which are dynamically influenced by the top-down feedback. This addition was inspired by biological studies in the brain, where there are bidirectional connections between lower-order and higher-order visual areas (Gilbert and Li 2013).

#### Visual Feature Extraction

Let *I* ∈ ℝ^H×W×3^ denote the input image. The visual system *V*(·) transforms the input into high-level visual features:

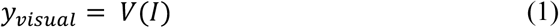

Here, *y_visual_* ∈ ℝ*^d^* represents the activation vector from the final visual layer (e.g., FC7), and *d* is the feature dimensionality.

### Emotion Recognition

The affective system, ***A***(·), receives the visual features in Eq. (1) and maps them to a set of 8 discrete emotion categories (e.g., amusement, fear, disgust, etc.) via three fully connected layers trained on emotion classification:

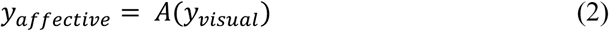

where **y*_affective_* ∈ ℝ^e^ and ***e*** = 8 is the number of emotion classes.

### Feedback Modulation

Following the long-range modulatory feedback framework proposed by (Konkle and Alvarez 2023a), we introduce feedback connections from the layer FC9 (source layer) in the affective system to the two target layers in the visual system: ℓ ∈ {*conv4, conv5*}. These connections modulate activations in the two visual layers using outputs from the affective layer FC9. Let ***X**_fc_* ∈ ℝ^B×F^ denote the batch-wise output of a fully connected layer (e.g., FC9) in the affective system, and let ***X**_conv_* ∈ ℝ^B×C×H×W^ be the activation tensor of a target convolutional layer in the visual system, with ***B*** as batch size, ***F*** as the number of units in the source layer, ***C*** as the number of channels, and ***H*** × ***W*** as spatial dimensions.

The feedback modulation proceeds as follows:

1. Normalization

The FC output is normalized using layer normalization:

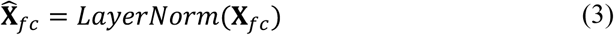

1. Activation Bounding

A tanh nonlinearity is applied to constrain the signal:

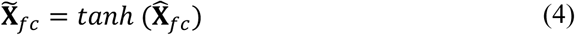

1. Learned Scaling

The bounded vector is scaled by a learnable weight vector ***s*** ∈ ℝ^F^, which encodes the contribution of each unit in the source layer:

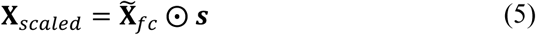

where ⨀ denotes elementwise multiplication.

1. Global Tuning Strength

An additional scalar parameter *λ* ∈ ℝ, referred to as the tuning strength, is applied to globally control the overall influence of the feedback signal:

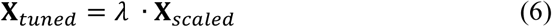

Unlike ***s***, which is learned during training, *λ* is manually controlled at test time to modulate the intensity of the top-down signal.

1. Reshaping and Broadcasting

**X***_turned_* is reshaped to shape (B, F, 1, 1) to prepare for channel alignment.

1. Channel Alignment via 1×1 Convolution

A 1×1 convolution projects the feedback signal to match the convolutional layer’s channel dimension C:

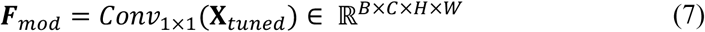

1. Multiplicative Modulation

The original activation **X***_conv_* is modulated through multiplicative gain control, consistent with biological evidence from attention studies (Treue and Trujillo 1999; Reynolds et al. 2000):

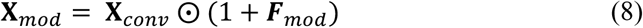

1. Nonlinearity

Finally, a ReLU activation is applied to the modulated output:

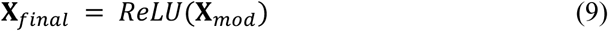

This mechanism allows the network to contextually modulate image processing in the visual system based on high-level affective representations, echoing theories of emotion-guided perception in biological systems (Pessoa 2008; Vuilleumier 2005; L. f. Barrett and Bar 2009; Pourtois et al. 2013).

#### Top-down Steering Modulation

Top-down steering modulation is designed to isolate the influence of category-level expectations on visual processing while holding the feedback circuitry fixed. In biological systems, top-down signals often convey abstract, task-relevant, or contextual information that biases sensory processing toward behaviorally relevant features ((Vuilleumier 2005; Reynolds and Heeger 2009; Summerfield and de Lange 2014).

It uses the same feedback pathways described earlier. The only difference is that instead of taking the FC9 activation (**X***_fc_*) from the current input image as the feedback signal, the modulation begins with a category prototype vector (*p*_k_ ∈ ℝ^F^) computed as the mean FC9 activation across validation exemplars of category *k*:

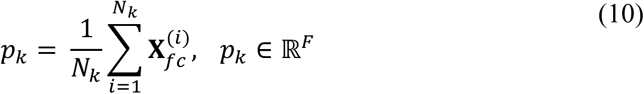

where *N*_K_ is the number of validation images in category *k*, and 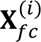 is the FC9 activation vector for image *i*. This prototype replaces **X***_fc_* at the normalization step, while all subsequent operations remain unchanged.

### fMRI Dataset

#### Participants

The experimental protocol was approved by the University of Florida Institutional Review Board. Twenty healthy volunteers with normal or corrected-to-normal vision participated in fMRI scanning after providing written informed consent (mean age = 20.4 ± 3.1 years; 10 male, 10 female).

#### Experimental Paradigm

Participants passively viewed affective images selected from the International Affective Picture System (IAPS; Bradley and Lang 2007) while simultaneous EEG-fMRI was acquired (EEG data are not analyzed in this study). Each picture was presented for 3 s, followed by a variable interstimulus interval (ISI) of either 2800 ms or 4300 ms. Jittered ISIs were employed to minimize temporal predictability and optimize event-related hemodynamic modeling. Each session consisted of 60 trials (one picture per trial), and participants completed five sessions in total. Across sessions, the 60 images were randomized in order. Stimuli were displayed on an MR-compatible monitor outside the bore and viewed through a reflective mirror. Participants were instructed to maintain central fixation throughout (see Fig. 8).

**Fig. 8.**
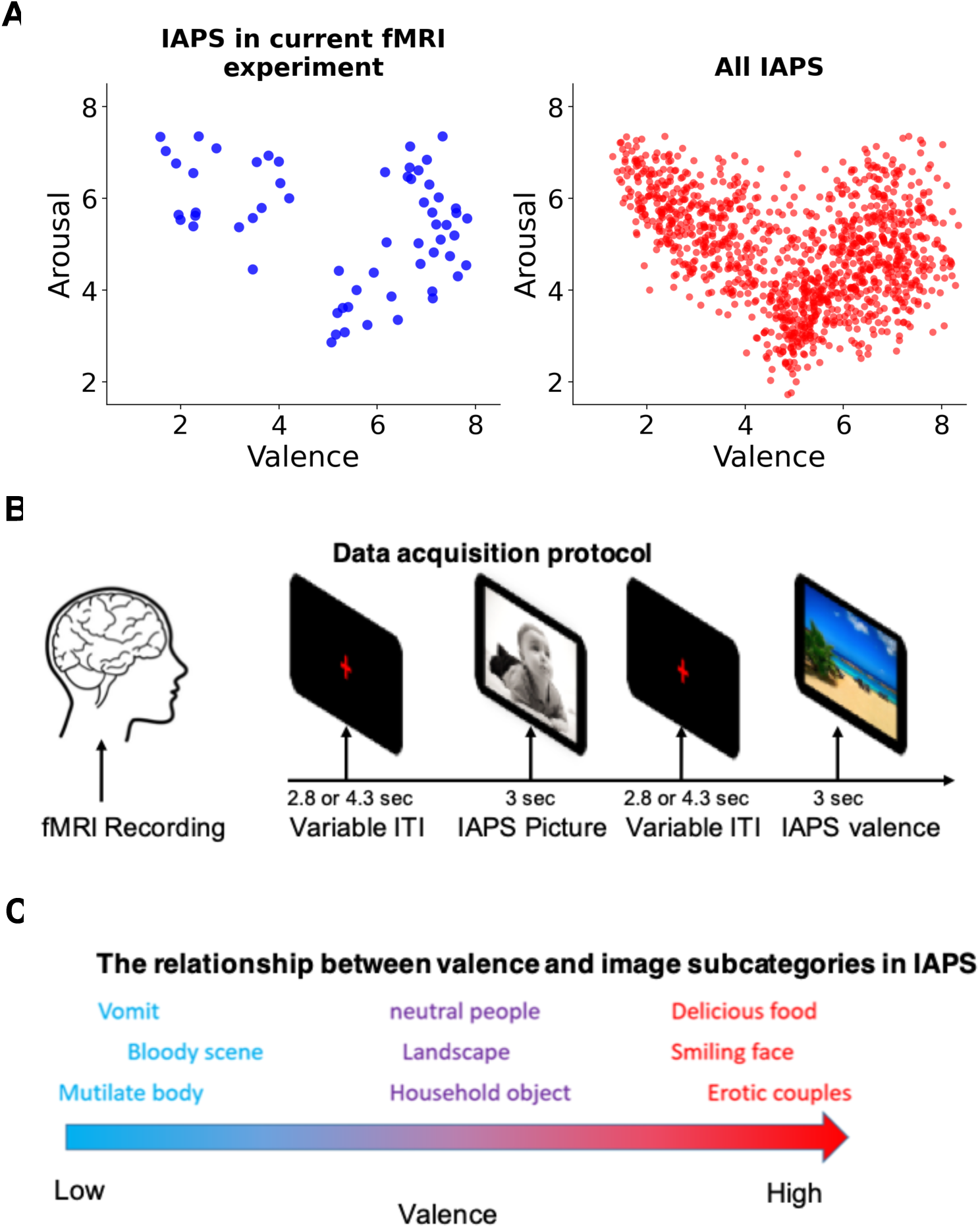
fMRI data acquisition and the distribution of the emotion variables in dataset IAPS. **A**. The protocol of brain data recording. It shows an image to a subject every 2.8 or 4.3 seconds. Forty-eight images are randomly selected from IAPS. The interval resting time is randomly set to be 2.8 or 4.3 seconds to reduce the priming rate before seeing an image. **B**. The emotion valence and arousal distribution in IAPS. (left) includes the 60 images, and (right) includes all IAPS images. As one can see, the selected 60 images are still following the same distribution pattern as all IAPS images. C. The relationship between valence and subcategories in IAPS images. Low valences are mostly related to vomit, bloody scenes, and mutilate body; the middle range valences are more associated with neutral people, landscapes, and household objects; the high valences include delicious food, smiling faces, and erotic couples. Example images shown in panel B are from the International Affective Picture System (IAPS; (Bradley and Lang 2007).

#### Data Acquisition and Preprocessing

Imaging was conducted on a 3T Philips Achieva scanner (Philips Medical Systems). Functional scans were acquired with the following parameters: TR = 1.98 s, TE = 30 ms, flip angle = 80°, 36 slices, FOV = 224 mm, matrix = 64 × 64, voxel size = 3.5 × 3.5 × 3.5 mm³. Slices were collected in ascending order and aligned parallel to the AC–PC plane. A high-resolution T1-weighted anatomical scan was also obtained (see Fig. 8).

Preprocessing was performed in SPM (http://www.fil.ion.ucl.ac.uk/spm/). The first five volumes from each run were discarded to allow for scanner equilibration. Standard steps included slice-timing correction, realignment (six motion parameters), normalization to the Montreal Neurological Institute (MNI) template with resampling to 3 × 3 × 3 mm³ voxels, and spatial smoothing with an 8 mm FWHM Gaussian kernel. A high-pass temporal filter (1/128 Hz) was applied to remove low-frequency drifts.

#### Single-Trial Beta-Series Estimation

To capture trial-specific responses, we applied the beta-series approach (Mumford et al. 2012). For each trial, a general linear model (GLM) was fit with the target trial modeled by its own regressor, while all remaining trials were grouped into a separate regressor. Six motion parameters were included as nuisance covariates. This process produced a unique beta estimate for every trial. For reliability, beta estimates corresponding to repeated presentations of the same picture across sessions were averaged, resulting in 60 distinct picture-specific activation patterns per participant. These trial-level patterns served as input for representational similarity analysis.

### Representational Similarity Analysis (RSA)

To evaluate how well the EmoFB network model captured category-level structure in its internal representations, we performed a representational similarity analysis comparing model-derived representational similarity matrices (RSMs) to a predefined theoretical RSM.

#### Theoretical RSM

The theoretical RSM encodes idealized representational geometry in which stimuli belonging to the same emotion category are maximally similar (similarity = 1), and stimuli from different categories are maximally dissimilar (similarity = 0). This matrix reflects the hypothesis that emotion category identity should shape the representational structure of the model.

#### Model-based RSMs

For each network layer (Conv1–Conv5, FC6–FC9), we extracted activation vectors in response to all test images and computed an empirical RSM by calculating Pearson correlations between all image pairs. This was done under two feedback conditions:

- Intrinsic Feedback: Top-down modulation was derived from instance-specific activations of the source layer (e.g., FC9) from the previous feedforward pass. This mechanism reflects internal recurrent processing within the model, where the feedback signal is stimulus-dependent and dynamically generated.
- External Steering: Top-down modulation was instead driven by a category-level prototype signal, computed as the average activation of the source layer (e.g., FC9) across all training exemplars belonging to the same emotion category. This form of feedback introduces an externally defined expectation that generalizes across instances.

Separate RSA analyses were conducted for each of the three task contexts: single image, side-by-side image, and overlay image.

#### Comparison to theoretical RSM

To quantify the alignment between empirical and theoretical representational structures, we computed the Pearson correlation between the lower triangular portions (excluding the diagonal) of each empirical RSM and the theoretical RSM. This resulted in a similarity score per layer, per task, and per feedback condition.

#### Statistical Analysis

Paired two-sided t-tests were used to compare the theoretical RSM similarity scores between the external steering and intrinsic feedback conditions at each layer, using asterisk notation: p < 0.05 (*), p < 0.01 (**), and p < 0.001 (***).

### Comparison with the human brain RSM

We performed representational similarity analyses to examine how closely the internal representations of the EmoFB model aligned with brain activity patterns across multiple visual and affective regions. This analysis was based on functional MRI (fMRI) data from 20 participants who viewed 60 emotional images selected from the IAPS. The image set was evenly divided into three affective categories**—**pleasant, neutral, and unpleasant, with 20 images per category. The same 60 images were used as input to the EmoFB network.

#### Brain RSM Construction

Neural activation patterns were extracted from six bilateral regions of interest: early visual areas (V1–V4), the lateral occipital complex (LOC), and the amygdala. For each image and subject, we extracted voxel-wise beta estimates from each ROI. The resulting *voxel × image × subject* matrices were cleaned by removing missing values (NaNs) and standardized using z-score normalization within each subject and image.

For each subject *s* and image *i*, we normalize the voxel activation ***v***_i,s_
∈ ℝ^v=^ as:

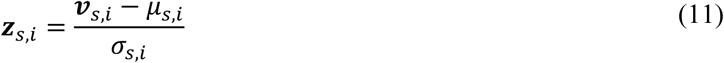

where ***μ****_s,i_* and ***σ****_s,i_* are the mean and standard deviation of the non-NaN voxel activations for subject *s* and image *i*. We then compute the average normalized activation across all subjects for each voxel and image:

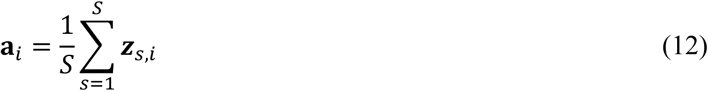

where ***a****_i_* ∈ ℝ^v=^ is the averaged activation vector for image *i*. For each pair of images (*i*, *j*), we compute the Pearson correlation:

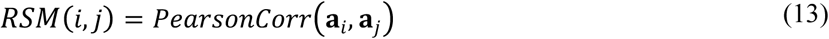

This yielded a 60 × 60 brain RSM for each ROI.

#### DNN RSM Construction

We extracted activation vectors from selected layers of the EmoFB model (Conv1–Conv5, FC6–FC9) in response to the same 60 IAPS images. RSMs were computed for each layer under two conditions:

- Before Steering: Activation patterns from the initial feedforward pass, prior to any top-down modulation.
- After Steering: Activation patterns from the final feedforward pass, following application of category-level external steering.

Pairwise Pearson correlations *r* between image activation vectors were computed, forming a 60 × 60 model RSM for each layer and condition.

#### Brain-Model Similarity

To assess representational correspondence, we computed the Pearson correlation between the lower triangular (off-diagonal) values of each DNN RSM and its corresponding brain RSM. This analysis was performed for each ROI and layer, both before and after steering.

#### Statistical Analysis

Similarity scores were compared using paired two-sided t-tests between the Before Steering and After Steering conditions. Significance levels were denoted as follows: p < 0.05 (*), p < 0.01 (**), p < 0.001(***), and “ns” for non-significant results.

### Top-down steering in advance of stimulus input

To test the extent to which abstract, category-level expectations can bias the initial feedforward sweep of visual processing, in the pure top-down steering condition, the EmoFB model received visual input through a single feedforward pass, whereas intermediate visual layers (e.g., Conv4 and Conv5) were already modulated by externally supplied category-level steering signals. In other words, the activity of the network was already biased by the steering signal in anticipation of the visual input. No recurrent passes or additional stimulus presentations were used, ensuring that any modulation originated solely from the top-down steering input.

Each test image was presented once to establish a baseline condition. In the pure external steering setting, the model did not reprocess the image or rely on recurrent passes. Instead, a category-level steering signal, computed as a prototype vector from validation set exemplars, was injected into designated layers during a single feedforward pass. The validation set was used to generate these prototypes to avoid overlap with training images and ensure an unbiased source of category-level information. This signal propagated forward through the network to generate an emotion prediction without direct image input. We evaluated two conditions:

- Pure Feedforward: A standard forward pass with the test image, with no external modulation.
- Pure External Steering: A forward pass using only the category-level steering signal to modulate internal representations.

We assessed Top-1 classification accuracy across three emotion recognition tasks—single image, side-by-side image, and overlay image. Paired two-sided t-tests confirmed statistical significance. Significance levels were denoted as follows: p < 0.05 (*), p < 0.01 (**), p < 0.001(***), and “ns” for non-significant results. To examine the effect of steering intensity, we varied a scalar tuning strength parameter (range: 0.0 to 3.0 in 0.5 increments), which scaled the external modulation signal. Across all tasks, accuracy followed an inverted-U profile, peaking at moderate strength (1.5). One-way repeated measures ANOVA revealed a significant main effect of tuning strength (*F* > 500, *p* < 1e−50), and post hoc tests showed significant pairwise differences between tuning levels (denoted by *, **, or ***). To evaluate whether pure top-down steering enhanced category structure, we applied the same representational similarity analysis used in Fig. 4, comparing each model layer’s RSM to a theoretical category-based RSM. Statistical differences between conditions were assessed using paired t-tests, where p < 0.05 (*), p < 0.01 (**), p < 0.001(***), and “ns” for non-significant results.

### Target-absent evaluation

Target-absent evaluation followed the same procedure as the pure top-down steering condition, with the only difference being the stimuli: test trials contained no image from the target category. As in the target-present case, a category-level steering signal (derived from validation exemplars) was injected into the designated layers during a single feedforward pass. This setup allowed us to quantify false alarm rates by measuring how often the model incorrectly reported the presence of the target emotion in the absence of a corresponding stimulus (i.e., false positives).

## Acknowledgements

This work was supported in part by the National Science Foundation grants 1908299 and 2318984 and the National Institutes of Health/National Institute of Mental Health grants MH112558 and MH125615, the University of Florida Artificial Intelligence Research Catalyst Fund, and the University of Florida Informatics Institute Graduate Student Fellowship. The funders had no role in study design, data collection and analysis, decision to publish, or preparation of the manuscript.

