## Supplemental Materials for "Visual Emotion Perception in a Deep Neural Network Model with Both Bottom-Up and Top-Down Connections"

Two topics related to the study reported in the main manuscript are addressed in this Supplementary Materials.

**Topic 1. Overview of all datasets used for training and testing the network**

Figure S1 provides a detailed overview of the EmoSet dataset splits and their use in training, validation, and testing.

Figure S1A shows the distribution of images across subsets. The training set contains between 8,434 (anger) and 15,837 (excitement) images per category. Validation sets range from 518 (anger) to 994 (amusement) images, while test sets range from 1,661 (disgust) to 3,014 (excitement) images. Although all categories are well represented, there is moderate class imbalance, with amusement and excitement containing the most samples, and anger and disgust containing the fewest. Figure S1B displays class weights derived from the training set to compensate for imbalance. For instance, fear and disgust receive the highest weight (~1.3) due to their smaller sample size, while amusement and excitement receive a lower weight (~0.7) given their larger sample count. Figure S1C shows the number of composite triplets constructed for the test set. Each emotion category contributes approximately 1,200 images per format (original, overlay, side-by-side), totaling ~3,600 per category. Counts per format range from 1,200 to 1,305, ensuring near-equal representation across conditions.

Together, these datasets form the basis for training, validating, and testing the proposed EmoFB network across diverse conditions.

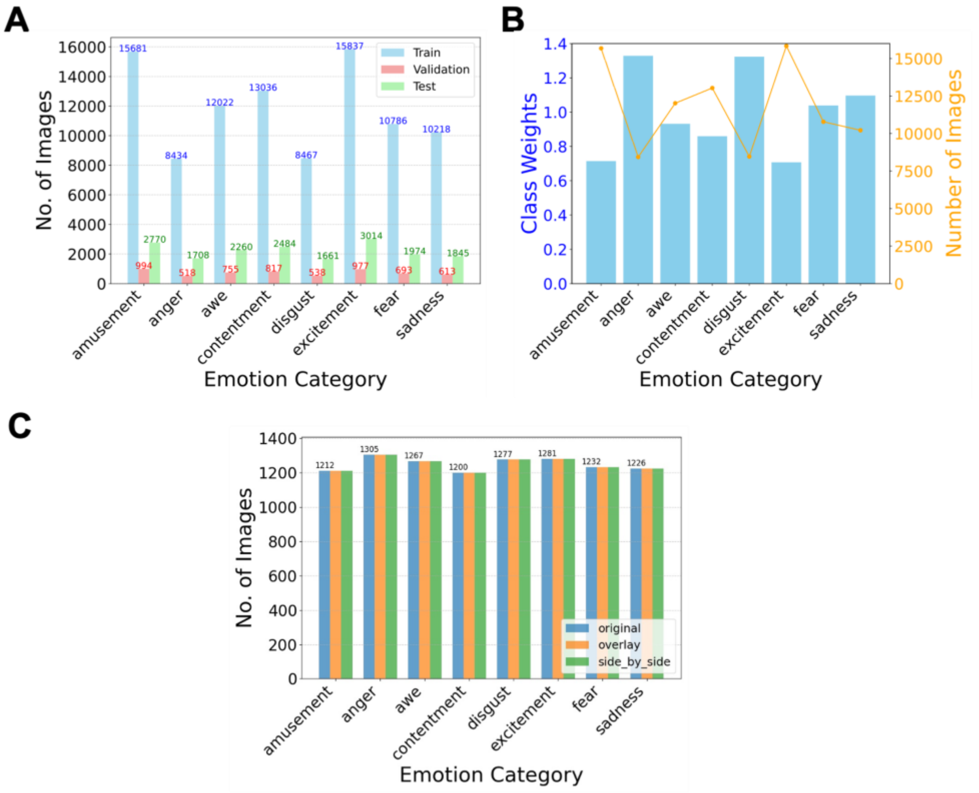


***Fig. S1 | Overview of all datasets used for training and testing the network. A.*** *Distribution of images across training, validation, and test sets in dataset EmoSet.* ***B.*** *Class weights derived from the training set of EmoSet to address class imbalance during network training.* ***C.*** *Number of composite triplets derived from EmoSet used for evaluating network performance during testing.*

**Topic 2. Noisy images and robustness of EmoFB with and without top-down steering**

We assessed the robustness of EmoFB under degraded visual conditions by adding Gaussian noise to test images (Figure S2). Noise level (NL) refers to the standard deviation of the Gaussian noise, with values ranging from 0.02 to 0.2.

Figure S2A shows sample inputs at increasing NL values (0.02, 0.05, 0.1, 0.15, 0.2). EmoFB processed these noisy inputs with one pure feedforward pass either with or without external steering applied. Figure S2B quantifies Top-1 accuracy across 10 independently trained models. At NL = 0.02, accuracy is ~70% in the no steering but rises to ~95% with steering. At higher noise levels, no steering feedforward accuracy declines steadily (~60% at NL = 0.05, ~50% at 0.1, ~30% at 0.15, and ~22% at 0.2), whereas steering maintains substantially higher performance (~95%, ~90%, ~75%, and ~60%, respectively). Across all tested NL values, steering significantly outperformed feedforward processing (***$p < 0.001$).

Overall, external steering improved recognition accuracy by 35–45 percentage points under noisy conditions, demonstrating that top-down modulation stabilizes performance and enhances robustness when sensory input is corrupted. This mirrors biological findings showing that feedback signals from higher-order areas can suppress noise and amplify relevant signals in sensory cortex. EmoFB's robustness to noise provides a computational parallel to how the brain leverages top-down feedback to maintain perceptual stability under degraded or ambiguous conditions.

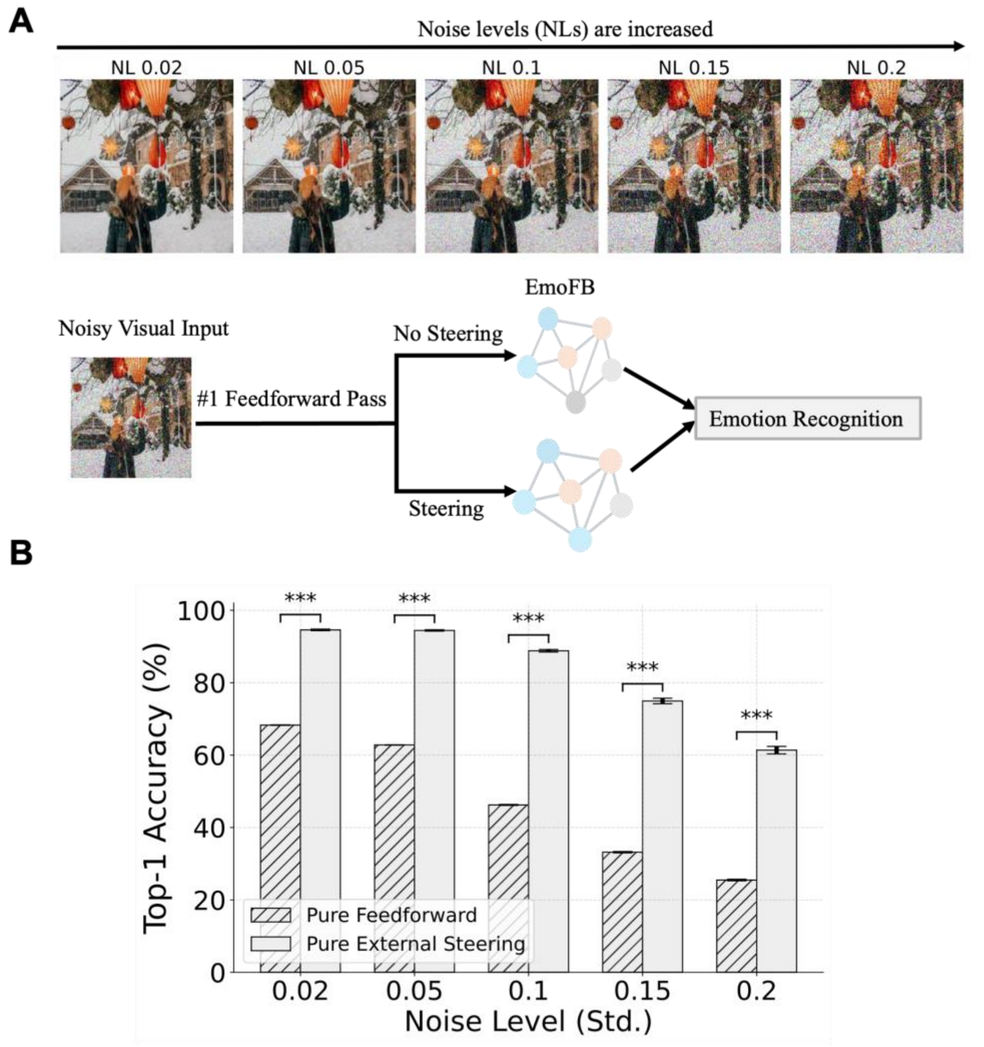


***S. Fig. 2 | Noisy image and model robustness with and without top-down steering.***

***A.*** *Sample visual inputs with increasing levels of Gaussian noise (NL = noise level standard deviation), ranging from 0.02 to 0.2. A schematic illustrates the EmoFB* *model's response to a noisy image during the first feedforward pass, comparing two processing ways: one without steering and one with top-down steering applied to the visual system.* ***B.*** *Top-1 emotion recognition accuracy of the EmoFB model under varying noise levels, comparing the pure feedforward condition (hatched bars) to the pure external steering condition (solid gray bars, tuning strength = 1.5). Steering significantly improved performance across all noise levels (**** $p < 0.001$*). Error bars denote standard error of the mean (SEM) across 10 independently trained models.*
